# Oxidative stress mediates cardiac electrophysiological injury in inhalation exposure to flavored vaping products

**DOI:** 10.1101/2025.05.31.657128

**Authors:** Obada J. Abou-Assali, Michelle Reiser, Mengmeng Chang, Bojjibabu Chidipi, Ryan Asswaytte, Yuting Zhang, Charles Szekeres, Laurent Calcul, Sami F. Noujaim

## Abstract

**Background:** Electronic nicotine delivery systems (ENDS) heat “E-liquids” to generate “E-vapor”, an aerosolized mixture containing nicotine and flavors. Flavored ENDS are popular among teens who vape, however, the possible cardiac electrophysiological harm of inhalation exposure to flavored ENDS are not fully understood.

**Objective:** To test if inhalation exposure to flavoring carbonyls in E-liquids compromises mitochondrial integrity, increases oxidative stress, and leads to cardiac electrophysiological toxicity.

**Methods:** Gas chromatography mass spectrometry (GC/MS), flow cytometry, oxygen consumption rate (OCR) measurement, in-vivo programmed electrical stimulation (PES), and multielectrode array (MEA) were used in atrial like HL-1 myocytes, hiPSC derived cardiomyocytes, and mice overexpressing mitochondrial catalase (mCAT).

**Results:** We compared the toxicity of E-vapor exposure from 30 differently flavored E-liquids in HL-1 cells using apoptotic annexin V flow cytometry. Most E-liquids were toxic. We identified the major flavoring carbonyls in these E-liquids and quantified their concentrations using GC/MS. Linear regression analysis showed that toxicity correlated with carbonyls concentration. Using flow cytometry of CellROX and TMRE staining, HL-1 cells exposed to flavored E-vapor showed increased reactive oxygen species and depolarized mitochondrial membrane potential. Additionally, exposure decreased OCR in these cells. In-vivo inhalation exposure to flavored E-vapor increased the inducible ventricular tachycardia duration in WT but not in mCAT mice compared to controls. MEA recordings in hiPSC derived cardiomyocytes exposed to flavored E-vapor, with or without nicotine, resulted in changes in the spontaneous beating rate.

**Conclusions:** Inhalation exposure to flavored ENDS negatively affects ventricular electrophysiology, in part via adverse mitochondrial remodeling, and increased oxidative stress.

## INTRODUCTION

Vaping, or the use of flavored electronic nicotine delivery systems (ENDs), remains a serious public health issue for school aged children and young adults.^1–4^ Recently, the CDC estimated that around 2.2 million middle and high school students reported vaping.^5^ It is also estimated that about 88% of these students specifically prefer to use flavored ENDS.^6^

Vaping is a form of electronic nicotine delivery where a coil is heated to aerosolize an “E-liquid”, a nicotine containing solution. E-liquids are a mixture of base humectants - propylene glycol and vegetable glycerin - with varying concentrations of flavoring constituents, and nicotine. The generated “E-vapor” is an inhalable smoke-like aerosolized mixture of nicotine, flavorings, humectant particles, and their resulting aldehyde byproducts.

It is becoming increasingly clear that vaping harms the cardiovascular system.^7–10^ For example, endothelial dysfunction due to ENDS use has been examined. The findings congruently report that vaping exposure causes vascular stiffness, increased endothelial microparticles, decreased levels of nitric oxide, and elevated levels of reactive oxygen species.^4,11–14^ Also, animal studies have suggested cardiac structural and electrophysiological changes such as ventricular remodeling, sympathetic predominance in heart rate variability, and increased arrhythmogenesis.^7,15–19^ In vaped mice, Oakes et, al., reported right ventricular hypertrophy with increased right ventricular free wall thickness, alongside increased RV systolic blood pressure. Us,^15^ and others,^18,20–22^ have shown increased sympathetic predominance in heart rate variability, increased arrhythmogenesis, action potential alternans, and ECG changes in mice with inhalation exposure to ENDS.

Studies in human users have shown that vaping increases the Tp-e/QT, an ECG index of ventricular repolarization which is associated with increased risk of sudden death.^16,23^ Furthermore, a case report and a retrospective analysis are pointing towards a possible association between recent vaping and sudden cardiac arrest.^24,25^ The retrospective analysis by Bains et. al, reported that in 6 patients (1% of the cases reviewed) sudden cardiac arrest or sudden death were temporally associated with vaping use.^24^ Four patients experienced sudden cardiac arrest with ventricular fibrillation within minutes to hours of vaping, and two patients experience sudden death associated with vaping induced respiratory distress. Interestingly, the mean age of these patients was 23 ± 5 years old, further raising concerns about the possible electrophysiological harm of vaping in young users.

The majority of studies have focused on the role of nicotine in mediating cardiac harm, but it has been suggested that other components of the E-liquids such as the humectants may negatively affect cardiac electrophysiology.^22^ Although the harm of flavorings in other organ systems such as liver, lungs, vascular, and immune systems have been investigated,^26–34^ the possible cardiac harm of flavoring carbonyls has not been extensively studied.

Even though flavored ENDS remain popular among young users, and preclinical and clinical studies are pointing to possible adverse cardiac effects associated with vaping, more mechanistic insights into how vaping impacts the electrophysiology of the heart are needed. Thus, our objective in this study is to investigate if flavorings contribute to cardiac harm, and if inhalation exposure to flavored ENDS leads to ventricular arrhythmogenesis through increased oxidative stress.

## MATERIALS AND METHODS

### HL1 Cell Culture

HL-1 cells (mouse atrial myocytes) were obtained from the laboratory of Dr. Claycomb (Louisiana State University) and cultured following the recommended protocol.^35^ Briefly, cells were grown in Claycomb medium (Sigma, St. Louis, MO) and supplemented with 10% FBS (Sigma, St. Louis, MO), 0.1 mM norepinephrine (Sigma, St. Louis, MO), 2 mM L-glutamine (Sigma, St. Louis, MO), and penicillin/streptomycin (100 U/mL / 100 m/mL) on tissue culture plates (Corning, Corning, NY), coated with fibronectin/gelatin (Sigma, St. Louis, MO).

### E-Liquids and E-Vapor Extracts

The flavored E-liquids used in this study are manufactured by USA Vape Labs (Huntington Beach, CA). The 30 differently flavored E-liquids were Menthol, American Patriot Tobacco, Cuban, Euro Gold, Macchiato, Vanilla Custard (VC), Triple Strawberry, Straw Lime, Berry Pom, Guava, Peach Lemon, Amazing Mango, Azul Berries, Naked Unicorn, Berry Lush, GoNanas, Green Blast, Maui Sun, Lava Flow, All Melon, Hawaiian POG (POG), Peachy Peach, Mango ICE, Lava Flow ICE, Very Cool, Brain Freeze, Apple Cooler, Hawaiian POG ICE, Polar Freeze, and Apple Jax (APJ, Epic Juice, Santa Ana, CA). These E-liquids are 70% vegetable glycerin / 30% propylene glycol (70VG/30PG) and are stated by the manufacturer to contain 6 mg/mL free-base nicotine. Base only (70VG/30PG, Sigma-Aldrich), and base plus 6 mg/mL free-base nicotine (Sigma-Aldrich) solutions were prepared in house. E-vapor extracts were generated as we did previously,^15^ and following the same puff topography as we have done previously.^15^ In short, E-vapor extracts were prepared fresh before each experiment by bubbling 10mL of fresh, untreated, culture media with 15 puffs of E-vapor generated from the different E-liquids, or base, or base plus nicotine. The resulting E-vapor extracts had a concentration of 1.5 puffs/mL, as we have done previously.^15^ For the air control media, the same protocol was followed with the vape device powered off thereby bubbling 10mL of fresh, untreated, culture media with room air for an equivalent amount of time.

### Apoptosis Flow Cytometry

Apoptosis was measured using the FITC annexin V staining assay (BD, Franklin Lakes, NJ). HL-1 cells were plated in 12-well plates and cultured to ∼70% confluency before being treated with either control media, or with one of the E-vapor extracts at 0.75 puffs/mL, as we have done previously.^15^ After 24hrs exposure to E-vapor extracts, cells in each well were detached with Accutase (approximately 8 mins), pelleted by centrifugation, washed with PBS, and stained with FITC annexin V according to the manufacturer’s recommendation. Cells were then pelleted by centrifugation, washed and resuspended in PBS. DAPI was added immediately before reading the samples on a BD LSRII Cytometer (using 488 nm and 405 nm lasers for the respective excitation of FITC annexin V and DAPI fluorescence, and 530/30 nm and 450/50 nm for their respective emissions). Data analysis was carried out using FlowJo software. As we did previously, the toxicity index was calculated as 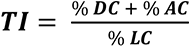 where %DC is the percentage of dead cells, %AC is the percentage of apoptotic cells, and %LC is the percentage of live cells per well.^15^

### Mass Spectrometry analysis of E-liquids

E-liquids were diluted 1:40 in pesticide grade methanol and ran through the Aglient 7890B gas chromatograph, mass spectrometer 5977B (GC-MS). The GC-MS parameters were optimized based on previous studies and the samples were run as we have done in our previously published work.^15^ MassHunter Workstation Qualitative Analysis Software (Version B.07.00 SP2) in conjunction with NIST MS Search 2017 Library was used for analysis. We then quantified the concentrations of the most abundant flavoring aldehydes (cinnamaldehyde, vanillin, and ethyl vanillin), and ketones (maltol and ethyl maltol), that we found in the screened E-liquids (19 out of 30 E-liquids). In order to quantify the concentrations of these carbonyls, we generated standard curves using 20, 50, 100, 150, 200 mM of each flavoring carbonyl in 70VG/30PG base. The total concentration of the flavoring carbonyls present in each of the E-liquid was then calculated. Figures were generated with OriginPro Lab Software 2018.

### Reactive Oxygen Species (ROS) Flow Cytometry

Oxidative stress was measured with the CellROX™ Green Flow Cytometry Assay Kit (Invitrogen; Catalog # C10492). HL-1 cells were cultured to ∼70% confluency before being treated with either control media or 0.75 puffs/mL Vanilla Custard E-vapor extract for 2hrs. Cells were then detached with Accutase (approximately 8 min), pelleted by centrifugation, washed and re-suspended in complete Claycomb medium. Control cells were pre-incubated with 5 mM N-acetylcysteine (NAC) for 30 mins, or with 200 µM tert-butyl hydroperoxide (TBHP) for 30 minutes. 500 nM CellROX Green was then added to all samples and incubated for 30 minutes at 37°C. Cells were then pelleted by centrifugation, washed and resuspended in PBS. Then, 5 µM SYTOX Red Dead Cell stain was added, and samples were read immediately on a BD LSRII Cytometer (using 488 nm and 639 nm lasers for the respective excitation of CellROX Green and SYTOX Red fluorescence, and 530/30 nm and 665/40 nm filters for the respective emissions).

### Mitochondria Membrane Potential (**ΔΨ**) Flow Cytometry

Mitochondria Membrane Potential was measured in HL-1 cells using tetramethylrhodamine ethyl ester (TMRE, Thermo Fisher; Catalog # T669). HL-1 cells were cultured to ∼70% confluency before being treated with either control media, 1 mM tert-butyl hydroperoxide (TBHP) as a positive control, or 0.75 puffs/mL Vanilla Custard E-vapor extract for 24hrs. Cells were then detached using 0.005% trypsin, pelleted by centrifugation, washed with PBS, stained with 50 nM TMRE and incubated for 20 minutes at 37 °C. Cells were then again pelleted by centrifugation, washed and resuspended in PBS, and DAPI was added immediately before reading the samples on a BD FACSCanto™ II Cytometer (using 488 nm and 405 nm lasers for the respective excitation of TMRE and DAPI fluorescence, and 585/42 nm and 450/50 nm for the respective emissions).

## OCR

The oxygen consumption rate (OCR) of HL-1 cells was measured with the XFp Cell Mito Stress Test by Agilent Seahorse. ATP-linked respiration was quantified with 1.5 µM oligomycin, maximal respiration with 1.5 µM phenylhydrazone (FCCP), and nonmitochondrial respiration driven by processes outside the mitochondria with 0.5 µM Rotenone and Antimycin. Two days prior to the experiment, 40,000 cells/well were plated in a gelatin/fibronectin precoated XFp mini cell culture plate (11.4 mm^2^ area) at 37°C. Cells were treated with either control media, or Apple Jax, or Hawaiian POG E-vapor extracts for 24hrs. Immediately before measuring, culture medium was replaced with the prewarmed assay medium (Agilent) containing 10 mM glucose, 1 mM pyruvate, and 200 mM L-Glutamate. Oligomycin, FCCP, and Rotenone/Antimycin were added to the cells in 20-minute intervals, according to the manufacturer’s instructions. At the end of the protocol, the culture media was gently removed, and 40 µl of RIPA lysis buffer was added, and cells were lysed using a 1 ml syringe. Protein concentrations were then quantified by BCA protein assay (Thermo Fisher Scientific) for normalization of the respiration parameters.

### Exposure of Animals to E-vapor

Exposure of mice to E-vapor was performed as we previously described,^15^ using the Smok Species Baby V2 (SMOKTech, Shenzhen, China) vaping device with the Baby V2 A2 dual sub coils (SMOKTech, Shenzhen, China) with a total resistance of 0.2 Ω, at 85 W. Mitochondrial catalase (mCAT) overexpressing mice (Jackson Laboratory, Bar Harbor, ME), and wild type (WT) littermates of both sexes were exposed to either room air or Vanilla Custard (VC) E-vapor containing 6mg/ml nicotine. For inhalation exposure, the animals were loaded into a custom-made chamber and were exposed to 4.7 second puffs of Vanilla Custard E-vapor at 1.4 L/ minute, every 2 minutes for a total of 60 puffs in a 2-hour period. After exposure, the animals were returned to their cages. Puff topography used a total puff volume of 110 mL,^15^ mimicking what has been described in human use.^36^ Mice were exposed 5 days a week, for a period of 5 weeks.

Control mice experienced the same handling as the vaped animals, were placed in a similarly modified chamber, in the same environment, where the only difference was that they were exposed to normal room air. Mice were individually housed in ventilated racks, with ad libitum access to food and water. All animal experiments were approved by the Institutional Animal Care and Use Committee at the University of South Florida.

### In Vivo VT Inducibility

Mice were anesthetized with 2% isoflurane, and a 1.2-French octapolar catheter (Millar, Houston, TX) was placed trans-venously into the right jugular vein and advanced into the right ventricle. Electrograms were recorded using the PowerLab platform (AD Instruments, Colorado Springs, CO). Programmed electrical stimulation (PES) for VT induction was performed by pacing the right ventricle at twice diastolic threshold with 1-s bursts from 20 to 50Hz, in 2-Hz increments as we did before.^15,37,38^

### hiPSC Derived Cardiomyocytes Culture and Extracellular Potential Recording

Human induced pluripotent stem cell derived cardiomyocytes, iCell^®^ Cardiomyocytes^2^, 01434 (Fujifilm; Catalog # R1220), were thawed, plated, and maintained according to MED64 Presto protocol for iCell^®^ Cardiomyocytes^2^. Cells were plated on the 24-well multiple electrode array (MEA) plates (24-well plate-eco, MED64; Parts # MED-Q2430M). The cells begin to beat spontaneously, periodically, and synchronously at around 4 days after plating. Cells remained beating robustly in culture for at least 14 days after thawing. hiPSC derived cardiomyocytes were treated with either control media, or with 0.15 puffs/mL of unflavored 70VG/30PG base without nicotine, or 0.15 puffs/mL Reconstituted Vanilla Custard E-vapor extract (RVC), or 0.15 puffs/mL Vanilla Custard without nicotine E-vapor extract (VC), or 0.15 puffs/mL Vanilla Custard with nicotine E-vapor extract (VC + Nic). Reconstituted Vanilla Custard E-liquid was made inhouse by supplementing unflavored 70VG/30PG base without nicotine with 79.2 mM vanillin, 48.8 mM ethyl vanillin, and 79.0 mM ethyl maltol to make RVC. This resulted in a reconstituted E-liquid with total carbonyls concentration of 207 mM, similarly to the original Vanilla Custard E-liquid. The Presto Multielectrode Array (MED64, Osaka, Japan) was used to simultaneously record the 16 unipolar extracellular potentials in each well of spontaneously beating hiPSC derived cardiomyocytes at 37°C and 95% O_2_ / 5% CO_2_. Wells that showed spontaneous, rhythmic activity before treatment were used. Five-minute recordings were obtained and quantified at baseline, and after treatment for 24hrs with the different E-vapor extracts or control media. The beating rate was calculated in the MEA Symphony analysis software (MED64, Osaka, Japan). Data for each electrode were then exported to verify the accuracy of the calculated beating rate.

### Statistics

Data are presented as average ± standard deviation. Student’s t-test and one-way analysis of variance (ANOVA) with post hoc Fisher’s LSD test on multiple comparisons were used as appropriate, and significance was taken at P < 0.05.

## RESULTS

We measured the toxicity of E-liquids in HL-1 mouse atrial cardiomyocytes using flow cytometry of FITC conjugated annexin V, an apoptotic marker, and DAPI for cell viability. Cells were cultured for 24hrs with control media or E-vapor extract for each of the thirty E-liquids, containing 6 mg/ml nicotine free base as per the label statement. Cells were then dissociated and stained with annexin V and DAPI. Flow cytometry analysis was performed to quantify the dead (high DAPI staining), apoptotic (high annexin V staining), and live (low annexin V and DAPI staining) cell populations, as we have previously done.^15^ The toxicity index of each of the E-liquid treated wells were calculated and normalized to air controls. Figure 1A shows a flow cytometry analysis of a control sample where the majority of cells are in the live quadrant. Panel B shows the flow cytometry analysis of cells treated with Macchiato E-vapor extract which caused a shift of cells to the apoptotic and necrotic quadrants. Figure 1C compares the normalized toxicity index after 24hrs treatment of HL-1 cells with 0.75 puffs/mL of the different E-vapor extracts. In Table 1 we grouped the E-liquids into seven types according to the description of the flavor profile as stated on the label. They were: 1-menthol, 2-tobacco, 3-vanilla, 4-simple fruits, 5-creamy fruits, 6-mixed fruits, and 7-creamy fruits and menthol. We also tested the unflavored E-liquid base containing 70VG/30PG and 6mg/ml nicotine. While some flavored E-liquids were more toxic than others, as seen in Figure 1C, the toxicities of all E-liquids including the unflavored base were significantly higher than that of air, except for Mango Ice and Amazing Mango.

**Figure 1:**
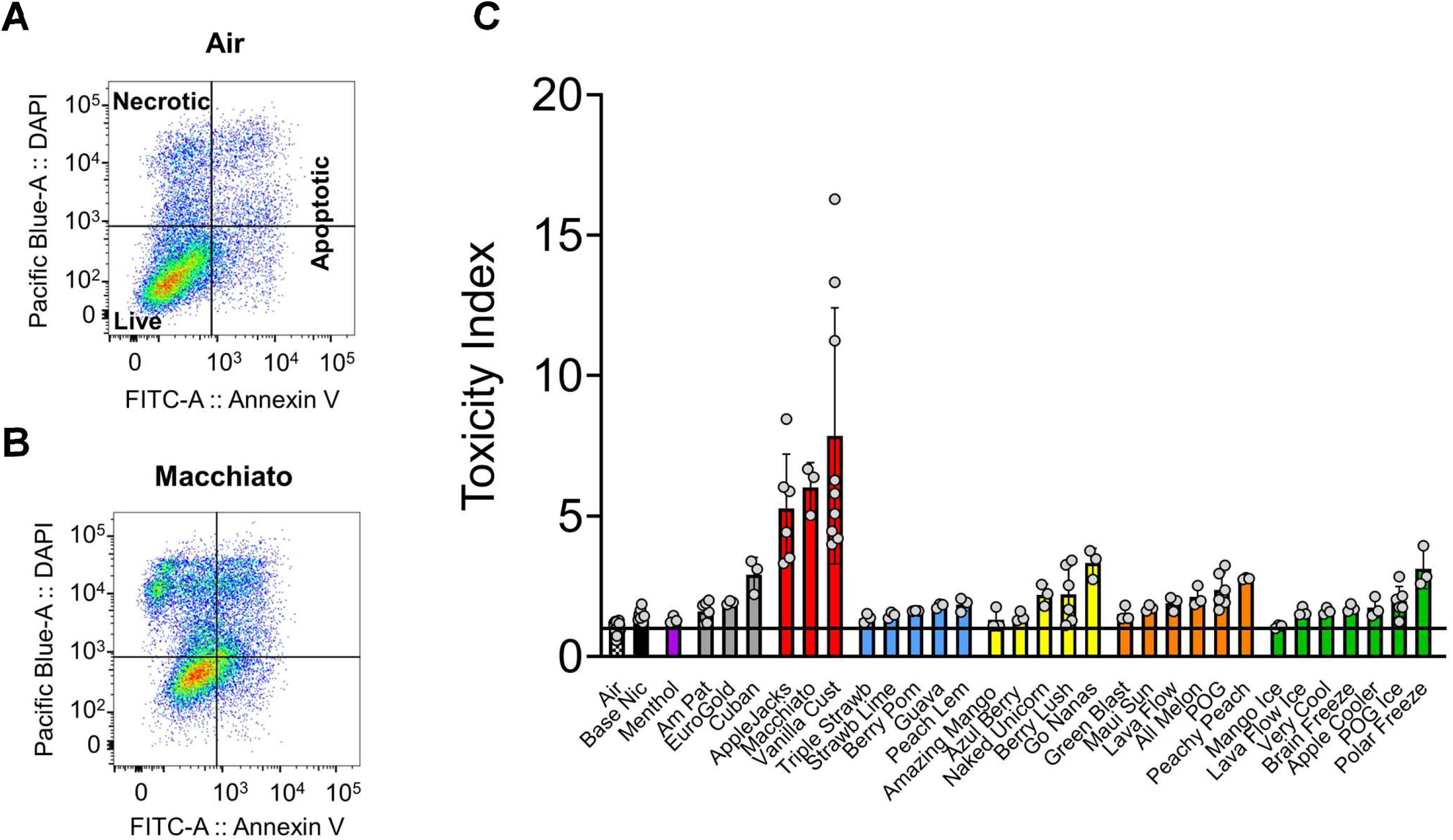
Toxicity of thirty differently flavored E-liquids in HL-1 cells cultured with 0.75 puffs/mL E-vapor extracts for 24hrs. Flow cytometry of annexin V and DAPI staining in HL-1 cells (Air control, **A**, and Macchiato, **B**). Live, apoptotic, and necrotic quadrants are indicated. **C:** Toxicity index of the E-liquids organized into 7 groups: 1-menthol (purple), 2-tobacco (silver), 3-vanilla (red), 4-simple fruits (blue), 5-creamy fruits (yellow), 6-mixed fruits (orange), and 7-creamy fruits and menthol (green). Unflavored base with nicotine (Base Nic) (black) and Air (black and white) were also included. All samples expect Amazing Mango and Mango Ice E-vapor extracts were found to be significantly toxic (p<0.05, each flavor vs its own air control, t-test, N= 4 to 6 each).

**Table 1:**
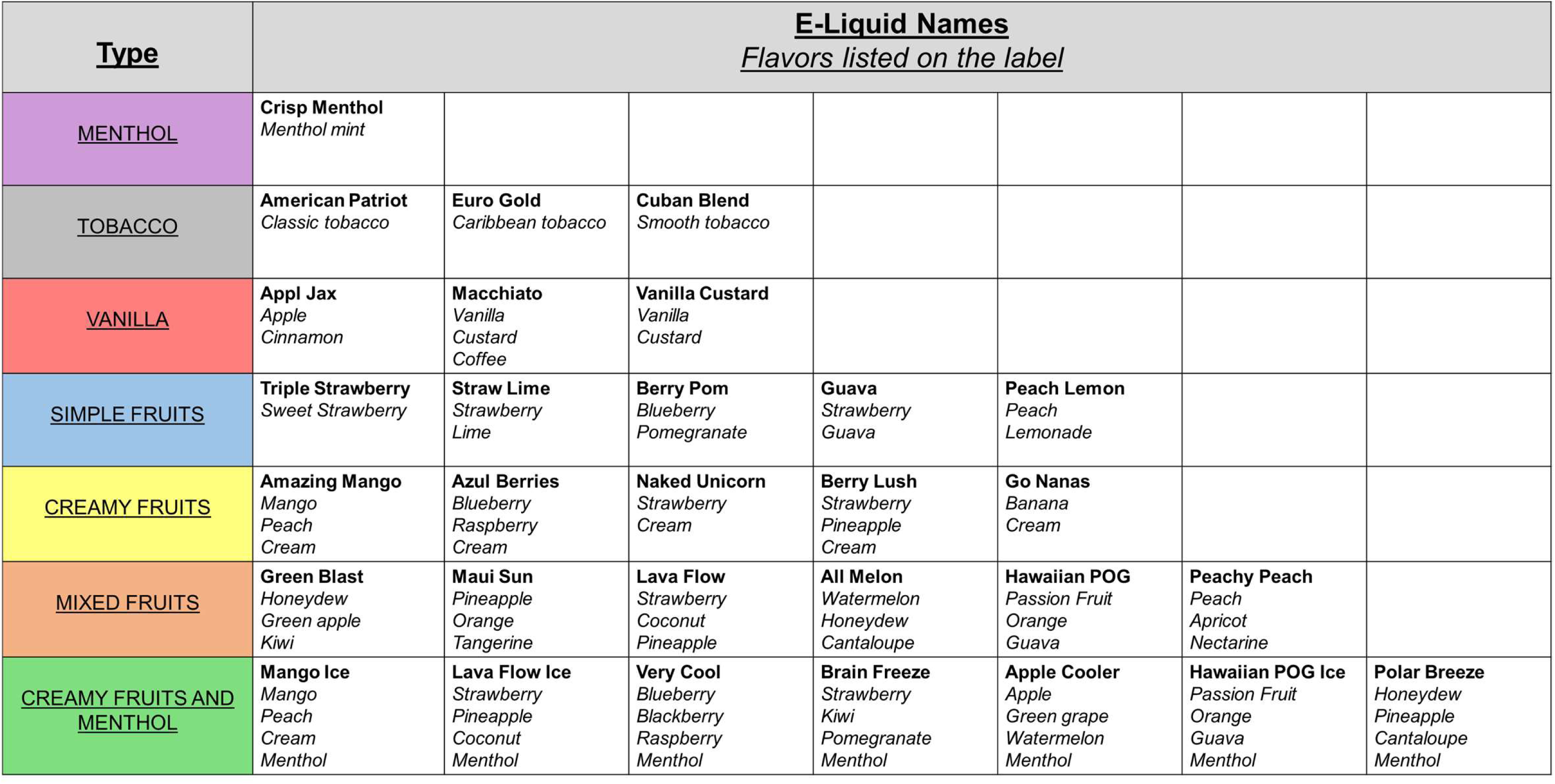
E-liquids grouped by type, with the E-liquids name (bold) and description of the flavor profile (italics) as printed on the manufacturer’s label.

We then used gas chromatography mass spectrometry (GC/MS) to analyze the chemical composition of the E-liquids listed in Table 1. Figure 2A shows the chromatograms of Macchiato (high toxicity index, Figure 1) E-liquid, and American Patriot Tobacco (low toxicity index, Figure 1) E-liquid. The propylene glycol peak was trimmed to enhance visibility of the smaller peaks of flavorings as customarily done.^39,40^ The glycerin and nicotine peaks are evident in addition to other flavorings peaks corresponding to flavoring carbonyl products such as the aldehydes vanillin and ethyl vanillin, and the ketone ethyl maltol. The peaks of vanillin, ethyl vanillin, and ethyl maltol are larger in the more toxic in the Macchiato E-liquid compared to the less toxic American Patriot Tobacco E-liquid (Figure 1). GC/MS analysis revealed that 19 of the 30 E-liquids contained vanilla (vanillin, and ethyl vanillin), cinnamon (cinnamaldehyde), or sweet (maltol, and ethyl maltol) flavoring carbonyls. In the rest of the E-liquids, other types of flavorings were present, but in small quantities. Since these carbonyls were the most abundant, we decided to focus on them. We then quantified using GC/MS the concentrations of the most abundant flavoring aldehydes. To quantify the concentrations, we constructed standard curves for each of the carbonyl compounds. Figure 2B is an example showing the GC/MS chromatograms of unflavored 70VG/30PG base that we spiked with known amounts of ethyl vanillin. Standard curves were constructed using the area under the curve. The concentration of the five carbonyls in the 19 E-liquids were then calculated. Some e-liquids contained one of the carbonyls (example, Strawberry Lime: 20.1mM ethyl vanillin, [total carbonyls]= 20.1mM, toxicity index= 1.48), while others contained a combination (example, Vanilla Custard: 79.2mM vanillin, 48.8mM ethyl vanillin, 79mM ethyl maltol, [total carbonyls]= 207mM, toxicity index= 7.85). Table 2 lists each of the 19 E-liquids, with their corresponding toxicity indexes from Figure 1, and the concentration of the different carbonyls in mM. Figure 2C is a plot of the total carbonyl concentration versus the toxicity index of the 19 e-liquids. Linear regression analysis revealed that toxicity correlated significantly with carbonyl concentration, where the coefficient of correlation was 0.84, p<0.001.

**Figure 2:**
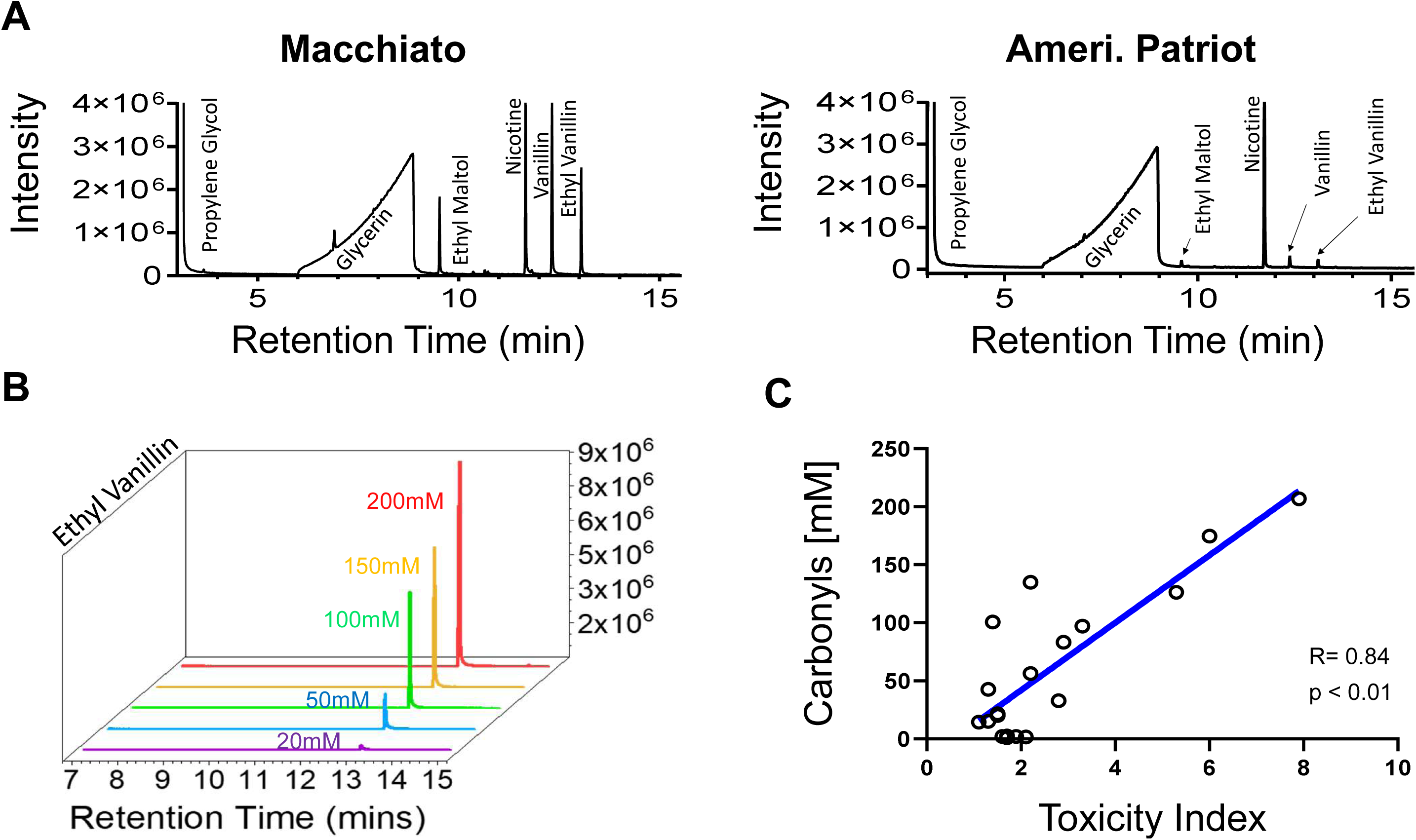
GC-MS analysis of flavoring carbonyls in the flavored E-liquids. **A:** GC-MS chromatograms of Macchiato (left) and American Patriot (right) flavored E-liquids. Glycerin, propylene glycol, and nicotine peaks are apparent in addition to peaks denoting flavoring carbonyls such as ethyl maltol, vanillin, and ethyl vanillin. **B:** Chromatograms of ethyl vanillin at different concentrations (20mM, 50mM, 100mM, 150mM, 200mM) used for constructing the standard curve, which was utilized to quantify ethyl vanillin concentrations in the different E-liquids. The same was done for vanillin, maltol, ethyl maltol, and cinnamaldehyde. **C:** Linear regression analysis of total carbonyl concentration versus toxicity index (R=0.84, p< 0.01).

**Table 2:** Toxicity index and carbonyl concentrations. The table lists the flavored E-liquid’s name, toxicity index, carbonyl concentration in mM for each of the five flavorings (vanillin, ethyl vanillin, maltol, ethyl maltol, and cinnamaldehyde), and the total carbonyl concentration.

| **<u>Flavor</u>** | **<u>Toxicity Index</u>** | **<u>Vanillin</u>** | **<u>Ethyl Vanillin</u>** | **<u>Maltol</u>** | **<u>Ethyl Maltol</u>** | **<u>Cinnamaldehyde</u>** | **<u>Total Carbonyls</u>** |
| --- | --- | --- | --- | --- | --- | --- | --- |
| Vanilla Custard | 7.9 | 79.2 | 48.8 |  | 79.0 |  | 207.0 |
| Macciato | 6.0 | 83.9 | 57.1 |  | 33.6 |  | 174.6 |
| Naked Unicorn | 2.2 | 46.7 | 57.4 | 2.2 | 28.5 |  | 134.8 |
| Apple Jax | 5.3 | 48.1 | 17.1 | 1.5 | 19.2 | 40.4 | 126.3 |
| Azul Berries | 1.4 | 33.3 | 44.2 |  | 23.3 |  | 100.8 |
| GoNanas | 3.3 | 34.2 | 22.1 |  | 40.8 |  | 97.1 |
| Cuban | 2.9 |  |  |  | 83.3 |  | 83.3 |
| Berry Lush | 2.2 | 17.2 | 28.6 | 2.4 | 8.2 |  | 56.4 |
| American Tobacco | 1.3 | 15.3 | 5.1 |  | 22.2 |  | 42.6 |
| Peachy Peach | 2.8 |  | 32.8 |  |  |  | 32.8 |
| GreenBlast | 1.5 | 17.5 |  | 1.5 | 2.6 |  | 21.6 |
| Strawlime | 1.5 |  | 20.1 |  |  |  | 20.1 |
| Mango | 1.3 | 12.6 | 2.6 |  |  |  | 15.2 |
| Mango ICE | 1.1 | 11.9 | 2.5 |  |  |  | 14.4 |
| BrainFreeze | 1.7 |  |  | 2.3 |  |  | 2.3 |
| LavaFlow | 1.9 |  |  | 2.0 |  |  | 2.0 |
| LavaFlow ICE | 1.6 |  |  | 2.0 |  |  | 2.0 |
| AllMelon | 2.1 |  |  | 1.6 |  |  | 1.6 |
| Applecooler | 1.7 |  |  | 1.0 |  |  | 1.0 |

Next, we measured reactive oxygen species (ROS) in HL-1 cells treated for 2 hours with 0.75 puffs/mL Vanilla Custard E-vapor extract to test whether vaping increases oxidative stress in cardiac myocytes. Figure 3A is the flow cytometry histogram analysis where Vanilla Custard (VC) treatment caused the cell population to shift toward higher ROS staining (red) compared to control (green). The black histogram shows that antioxidant pretreatment with 5 mM N-acetylcysteine (NAC) for 30 minutes blunted the effects of VC E-vapor extract on ROS generation.

**Figure 3:**
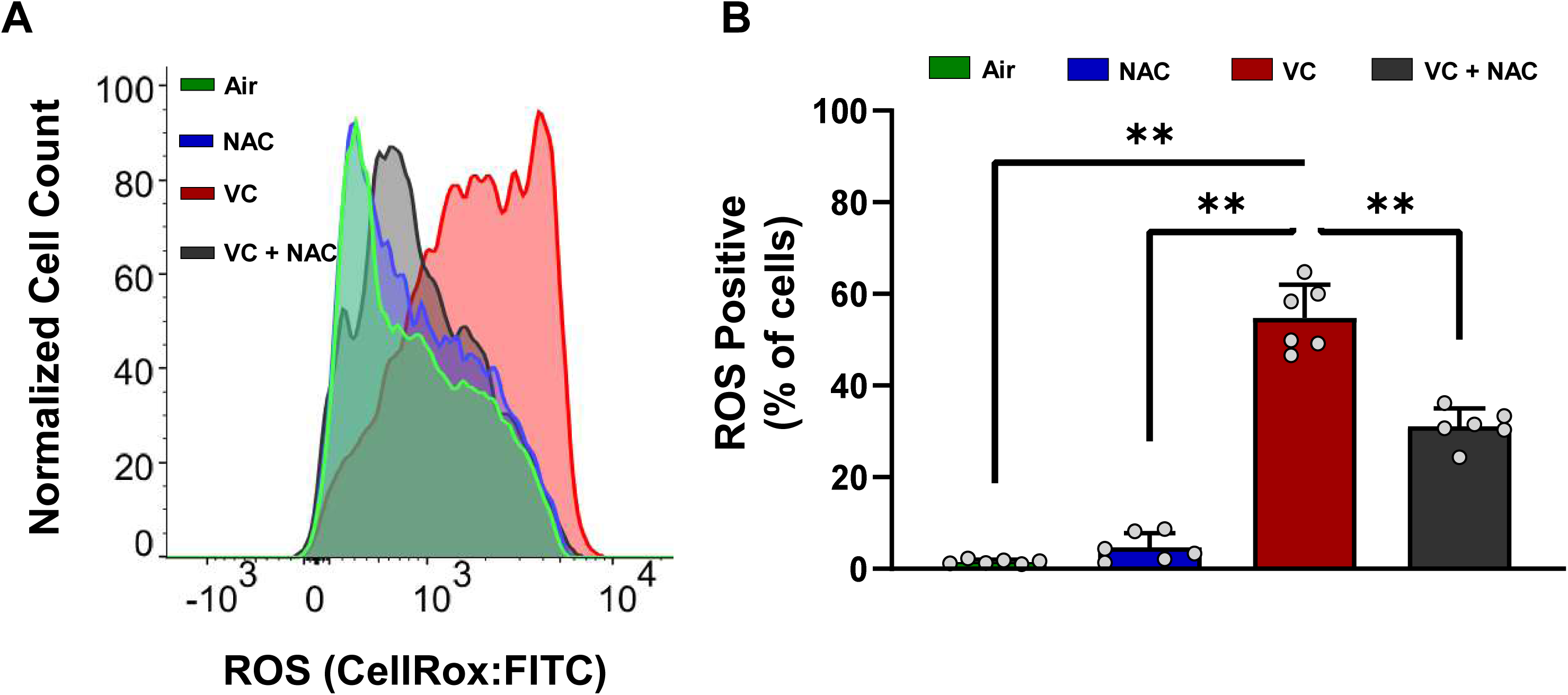
Quantification of CellRox staining in HL-1 cells with flow cytometry. **A:** Histogram of normalized cell counts for air control (green curve), 5 mM N-acetylcysteine pretreatment (NAC, blue curve), 0.75 puffs/mL Vanilla Custard E-vapor extract (VC, red curve), and 0.75 puffs/mL Vanilla Custard E-vapor extract with 5 mM N-acetylcysteine pretreatment (VC + NAC, black curve). **B:** Quantification of the % of cells with high ROS staining (**p<0.01, vanilla custard versus all other groups, one-way ANOVA with Fisher’s LSD test).

Blue represents control cells treated only with 5 mM NAC for 30 minutes. Panel B is a quantification of the percentage of cells with high ROS staining. A significant increase in ROS positive cells were observed in the VC treatment, and the NAC pretreatment significantly reduced ROS generation.

We also measured the mitochondrial membrane potential (ΔΨ) in HL-1 cells treated with either control media, TBHP, or VC E-vapor extract. Figure 4 panel A shows the flow cytometry histogram analysis where VC treatment (red) caused the cell population to shift toward lower TMRE fluorescence, suggesting a depolarization of the ΔΨ compared to control (green). Similarly, TBHP treatment (positive control, black) caused a depolarization of the ΔΨ. Figure 4B is the quantification of the percentage of cells with depolarized ΔΨ, which was significantly lower in control versus TBHP and VC treatments.

**Figure 4:**
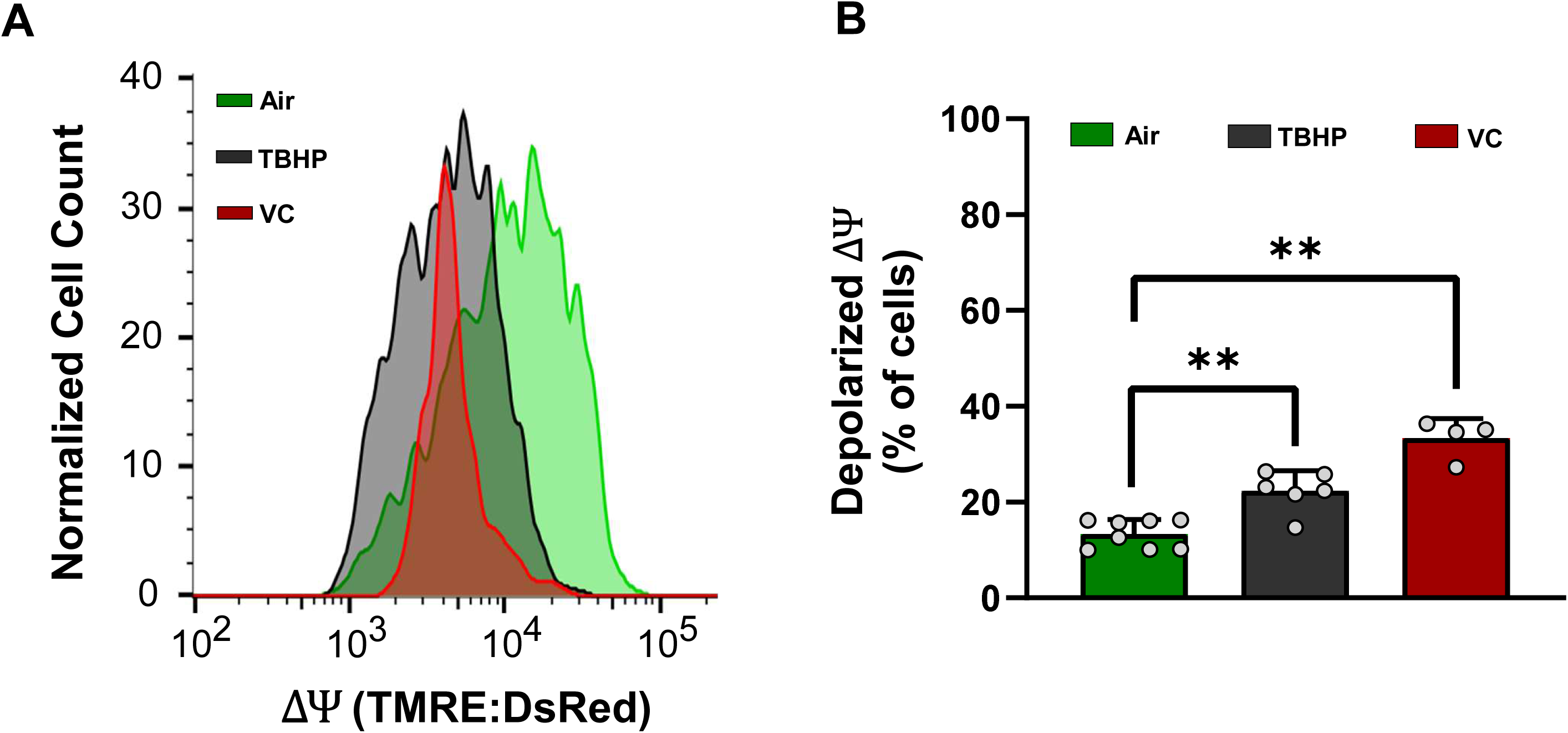
Quantification of TMRE staining in HL-1 cells with flow cytometry. **A:** Histogram of normalized cell counts for air control (green curve), 0.75 puffs/mL Vanilla Custard E-vapor extract (VC, red curve), and 1 mM tert-butyl hydroperoxide (TBHP, black curve). **B:** Quantification of the % of cells with low TMRE staining (**p<0.01, one-way ANOVA with Fisher’s LSD test).

We then compared cellular respiration in HL-1 cells after 24h treatment with 0.375 puffs/mL Evapor extract from an E-liquid with a higher toxicity (Apple Jax, APJ, toxicity index= 5.3) or lower toxicity (POG, toxicity index=2.3) as reported in Figure 1. Figure 5A is a graph of OCR versus time for control, APJ, or POG E-vapor extract treated cells. Figure 5B quantifies basal OCR (OCR before addition of Oligo), ATP linked OCR (OCR before Oligo minus OCR before FCCP), and maximal OCR (OCR before A/R minus OCR after A/R). Basal and ATP linked OCR were significantly depressed with APJ and POG treatment versus air, but maximal OCR was significantly lower in air versus APJ only.

**Figure 5:**
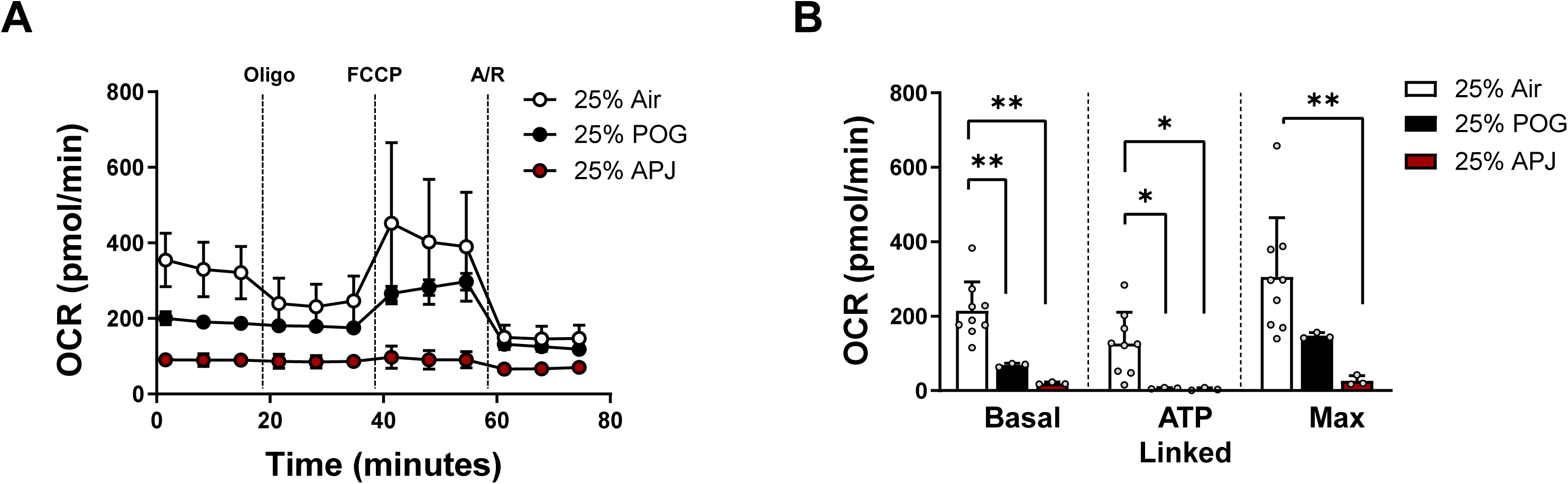
Oxygen consumption rate in HL-1 cells cultured with APJ and POG E-vapor extracts. **A:** Normalized OCR versus time in cells cultured for 24hrs with control media, or 0.375 puffs/mL APJ, or POG E-vapor extracts. The lines denote injections of Oligo (oligomycin), FCCP (phenylhydrazone), and A/R (Antimycin and Rotenone). **B:** Quantification of basal, ATP linked, and maximal OCR (*p<0.05, **p<0.01, one-way ANOVA with Fisher’s LSD test).

To investigate the role of inhalation exposure to flavored E-vapor on cardiac electrophysiology and examine the possible involvement of ROS, we used mitochondrial catalase (mCAT) overexpressing mice and their wild type (WT) littermates. mCAT overexpressing and WT littermate mice were vaped with Vanilla Custard E-liquid or with room air for 5 weeks. In vivo programmed electrical stimulation was performed. Figure 6A shows example ECG traces of inducible ventricular tachycardia (VT) in mice from the different groups. In Panel B we quantify VT duration.

**Figure 6:**
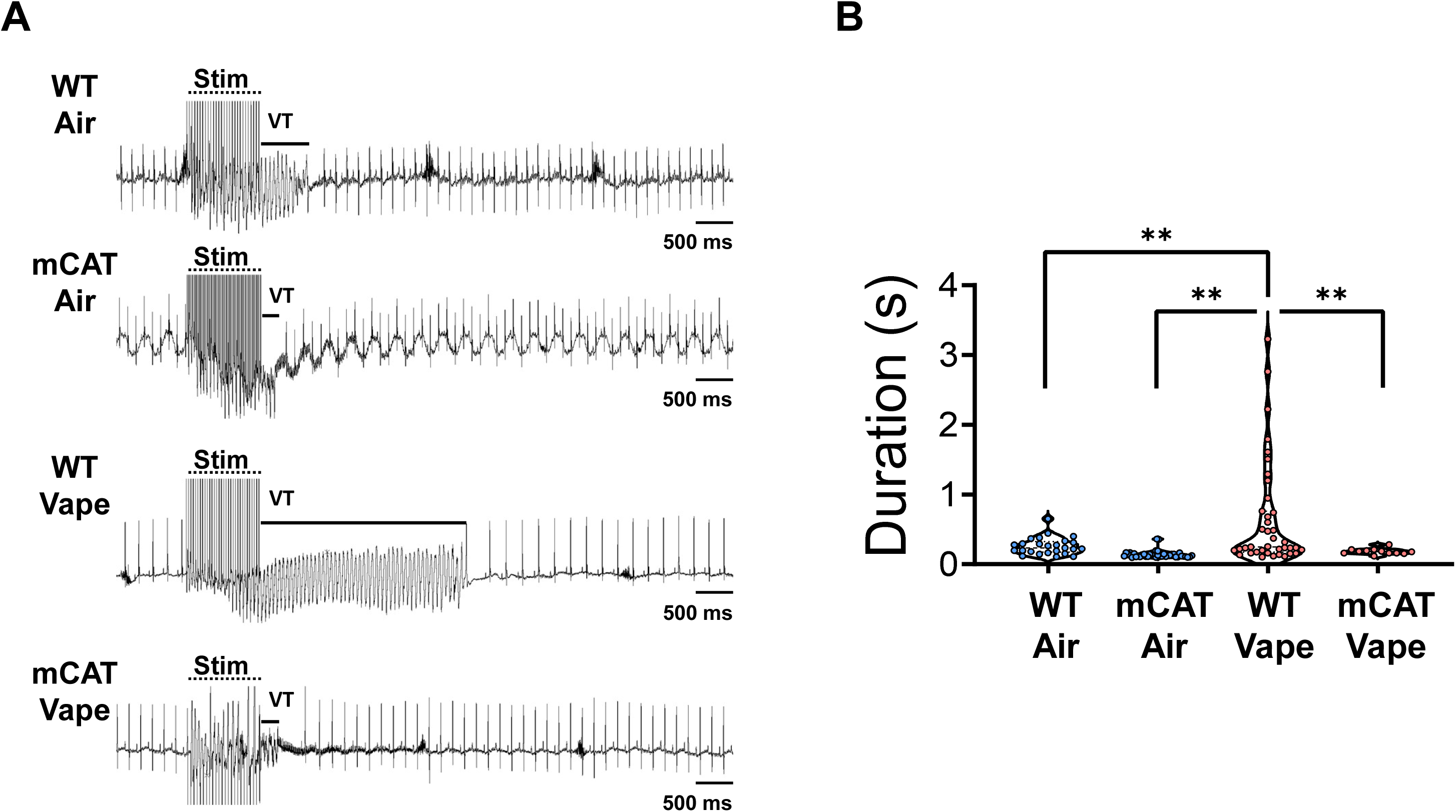
In vivo ventricular tachycardia (VT) inducibility in mCAT overexpressing and WT littermates exposed to room air or Vanilla Custard E-vapor. **A:** ECG traces of inducible VT in WT air, mCAT air, WT vape, and mCAT vape mice. **B:** Quantification of VT duration in the different groups (**p<0.01, one-way ANOVA with Fisher’s LSD test). Stim: burst pacing stimulation.

While the majority of animals were inducible, the VT duration was significantly longer in the vaped WT mice compared to all other groups. Importantly, overexpression of mCAT abrogated the effects of inhalation exposure to Vanilla Custard E-vapor on VT duration.

We then investigated whether exposure to flavorings alone, without nicotine, has an impact on cardiomyocytes. We tested in spontaneously beating hiPSC derived cardiomyocytes, the effects on beating rate of: 1-control media, 2-E-vapor extract of unflavored base without nicotine, 3-E-vapor extract of reconstituted Vanilla Custard (RVC), 4-E-vapor extract of Vanilla Custard without nicotine purchased from USA Vape Labs (VC), and 5-E-vapor extract of Vanilla Custard with nicotine (VC + Nic). We reconstituted Vanilla Custard E-liquid based on the concentrations of flavoring carbonyls determined by GC/MS and reported in Table 2. To do so, we used unflavored 70VG/30PG base without nicotine, supplemented with 79.2 mM vanillin, 48.8 mM ethyl vanillin, and 79 mM ethyl maltol to make the RVC E-liquid with a total carbonyl concentration of 207 mM. Figure 7A shows the electrograms of the same electrode recorded before and after 24hrs treatment for each group. Base, RVC, and VC accelerated the beating rate, whereas VC + Nic caused a deceleration. Panel B is the average beats per minute (BPM) before and after 24hrs treatment. The beating rate significantly changed after 24hrs treatment with Base, RVC, VC, and VC + Nic compared to pre-treatment baselines. Panel C shows the percent change in beating rate where RVC and VC resulted in significant acceleration, and VC + Nic resulted in significant deceleration.

**Figure 7:**
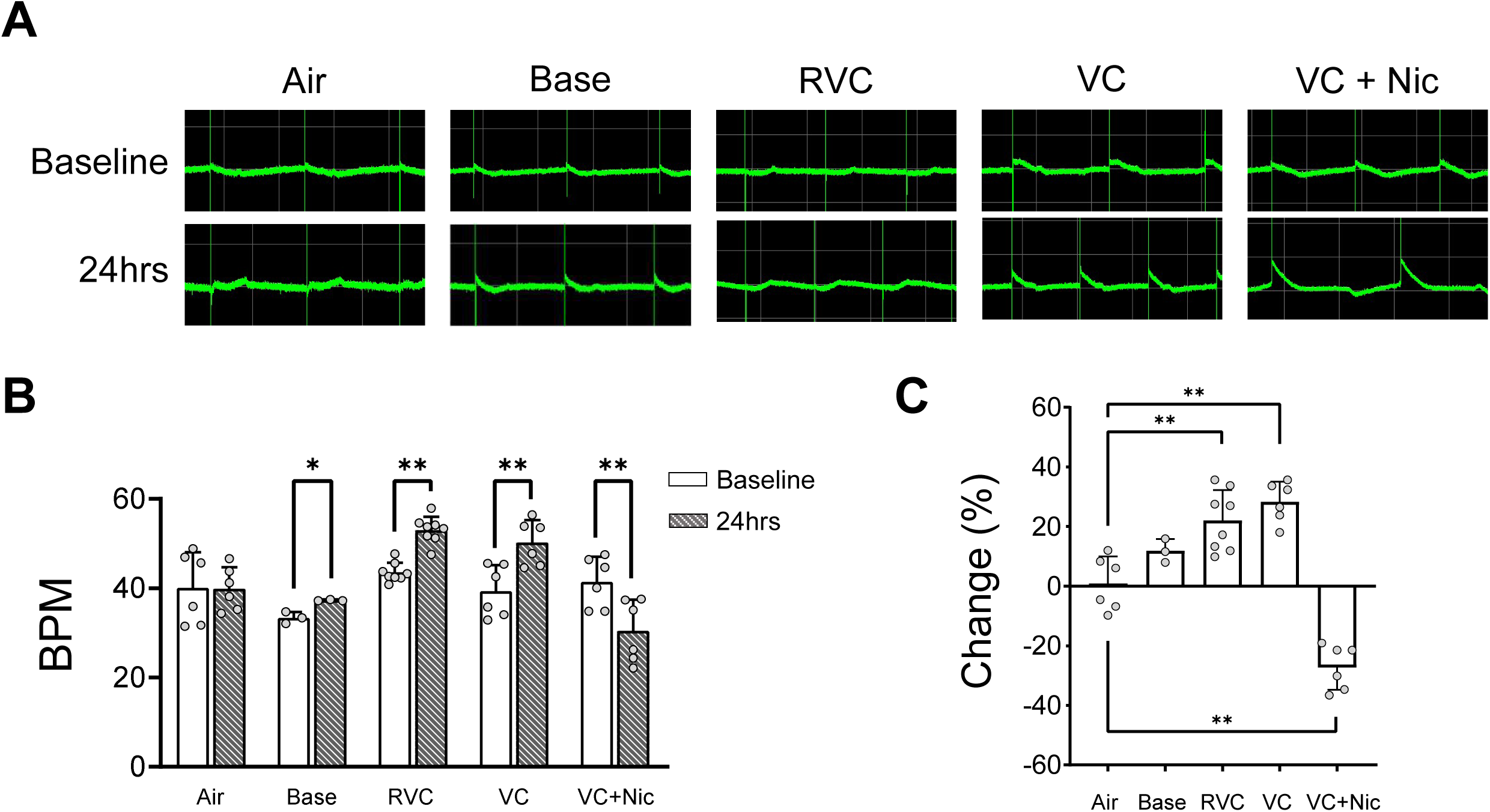
Beating rate analysis in hiPSC derived cardiomyocytes treated with either control media, or with 0.15 puffs/mL unflavored 70VG/30PG base without nicotine, or 0.15 puffs/mL Reconstituted Vanilla Custard E-vapor extract (RVC), or 0.15 puffs/mL Vanilla Custard without nicotine E-vapor extract (VC), or 0.15 puffs/mL Vanilla Custard with nicotine E-vapor extract (VC + Nic). **A:** Extracellular electrogram recordings before and after the different 24hrs treatments. **B:** Quantification of beating rate at baseline and after 24hrs treatment. (*p< 0.05, **p< 0.01, paired t-test). **C:** Quantification of percent change in beating rate (**p< 0.01 one-way ANOVA with Fisher’s LSD test).

## DISCUSSION

Our study suggests that inhalation exposure to flavored ENDS negatively affects ventricular electrophysiology, in part via mitochondrial dysfunction and increased oxidative stress. Additionally, our results indicate that exposure to flavoring carbonyls may exert toxicity on cardiac myocytes. Furthermore, oxidative stress and mitochondrial dysfunction may mediate increased ventricular arrhythmogenicity in inhalation exposure to flavored ENDS.

In this paper we investigated the toxicity of 30 differently flavored E-liquids on HL-1 cardiomyocytes. Most of the E-liquids tested (28 out of 30) were significantly toxic with some E-liquids being more toxic than others (Figure 1). We surmised that this was independent of nicotine content since all tested E-liquids contained 6 mg/mL nicotine as stated by the manufacturer. In line with our study, reduced cellular viability as a result of exposure to flavored ENDS has been reported by others, however the tests were conducted in non-cardiac cells.^39,41–43^ Our results also suggest that the flavoring constituents in the 30 different E-liquids drive the variation in toxicity. GC-MS quantification of carbonyls in the flavored E-liquids identified five major flavoring carbonyls (vanillin, ethyl vanillin, cinnamaldehyde, maltol, and ethyl maltol), and that toxicity correlated with the concentration of the total flavoring carbonyls (Figure 2). Sassano et. al. performed a high throughput assay of 148 E-liquids and found that vanillin and cinnamaldehyde correlated with toxicity in non-cardiac cells.^39^ Interestingly, E-liquids in the vanilla group were found to be more toxic than any other group and were found to have the highest concentrations of vanillin and cinnamaldehyde.

Since the accumulation of toxic carbonyls has been shown to cause an increase in oxidative stress,^44–46^ we investigated whether exposure to E-liquids increases ROS in cardiac myocytes. In HL-1 cardiomyocytes exposed to flavored E-vapor extracts, ROS significantly increased but was attenuated by pre-treatment with antioxidant NAC (Figure 3).^47,48^ We also assessed the effects of exposure on mitochondrial function by measuring ΔΨ (Figure 4) and OCR (Figure 5).

Vanilla Custard E-vapor extract depolarized the ΔΨ, and Apple Jax (a high toxicity E-liquid, Figure 1) impaired the cellular respiratory process more than POG (a low toxicity E-liquid, Figure 1) or air control. ROS is regularly produced as a consequence of the mitochondria’s normal cellular respiratory processes, and the high ROS accumulation that we observed may be in part due to flavored ENDS exposure mediated mitochondrial dysfunction. To further corroborate this contention, we tested in vivo the potential arrhythmogenicity of inhalation exposure to a flavored Eliquid and the possible role of mitochondria therein. mCAT overexpressing mice and their WT littermates were exposed to Vanilla Custard E-vapor or room air. In PES studies, WT mice exposed to Vanilla Custard E-vapor had significantly more sustained VT compared to the mCAT overexpressing animals (Figure 6). This suggests that the overexpression of mitochondrial catalase attenuated the effect of inhalation exposure on ventricular arrhythmogenesis and further points to the involvement of mitochondrial pathways in cardiac electrophysiological injury due to flavored ENDS exposure.

While nicotine’s harmful effects on the human heart have been known and well documented,^49–52^ it remains unclear whether the other components of flavored ENDS can also harm the heart.

Our experiments in hiPSC derived cardiomyocytes suggest that the base humectants alter the electrophysiology by accelerating the beating rate, and flavorings amplifies this effect. The presence of nicotine had the opposite effect of decelerating the spontaneous beating rate. It is possible that the effect of flavorings combined with those of cholinergic nicotinic receptors that have been recently described in ventricular myocytes contribute to this effect.^53^ However, further investigations are necessary to test this.

## LIMITATIONS

Our in vitro exposure studies do not accurately represent the actual plasma concentrations of the flavoring constituents, base humectants, nicotine, their thermal degradation byproducts, or blood metabolites. This is an important limitation that warrants improvements driven by a better understanding of these byproducts and metabolites concentration in the blood of ENDS users. It remains unclear if the ROS byproducts that have been identified in E-vapor initiate cellular oxidative stress or whether the components and thermal degradation byproducts in E-vapor directly cause oxidative stress. It is also possible that a combination of these scenarios results in ROS begetting ROS, leading to a cascade of amplification. Our results in vivo remain limited by the translatability of findings from mice models to humans. Pre-clinical and clinical studies are pointing to harmful consequences of ENDS exposure on the heart.^16,23–25^ Extrapolation of our findings in the mouse to the human should be done cautiously.

## ABBREVIATIONS

ANOVA: one-way analysis of variance APJ – Apple Jax E-liquid
BPM: Beats per minute
ENDS: Electronic Nicotine Delivery Systems FCCP – Phenylhydrazone
GC/MS: Gas Chromatography / Mass Spectroscopy mCAT – Mitochondrial catalase overexpressing mice MEA – Multielectrode array
NAC: N-acetylcysteine
OCR: Oxygen consumption rate
PES: Programmed Electrical Stimulation POG – Hawaiian POG E-liquid
ROS: Reactive oxygen species
RVC: Reconstituted Vanilla Custard E-liquid TBHP – Tert-butyl hydroperoxide
TMRE: Tetramethylrhodamine ethyl ester VC – Vanilla Custard E-liquid
VC + Nic: Vanilla Custard E-liquid with nicotine VT – Ventricular Tachycardia
WT: Wild Type

## AKNOWLEDGEMENTS

This study was funded in part by NIH R01ES032099 and AHA 24TPA1304716 grants to SFN.

